# Neuronal integrity is altered in the ipsilesional hand sensory territory in late stroke

**DOI:** 10.64898/2026.01.20.700729

**Authors:** Rafay Cheema, Nawal Cheema, Andrew Apostol, Carmen M. Cirstea

**Affiliations:** Boston University, Boston, MA; Physical Medicine and Rehabilitation, School of Medicine, University of Missouri, Columbia, MO

**Keywords:** somatosensory cortex, stroke, MR Spectroscopy, hand impairment

## Abstract

**Background:** Sensorimotor remapping plays a crucial role in the rehabilitation of hand function after stroke. While motor remapping has been intensively investigated at various levels, from functional to metabolic, limited attention has been given to somatosensory cortical remapping. This study extends our previous work showing functionally relevant metabolic alterations in radiologically normal-appearing or spared motor cortices, predominantly in the ipsilesional (stroke-injured) hemisphere, during the chronic phase of stroke. Precisely, we investigated the metabolic status of the ipsilesional primary somatosensory cortex (S1) and its relation to hand impairment in late stroke.

**Methods:** Fourteen individuals with a chronic (mean ± SD, 21.1 ± 36.9 months post-onset) ischemic subcortical stroke, exhibiting moderate hand impairment (Fugl-Meyer Upper Extremity, 44.8 ± 20.9; Jamar dynamometer, 54.1% ± 40.9% of the non-paretic hand), but without sensory deficits, and ten matched healthy controls participated. MR Spectroscopy markers of neuronal status (N-acetylaspartate) and neuronal-glial glutamatergic cycle/neurotransmission (glutamate-glutamine complex) were measured in the ipsilesional S1 (3T, Allegra Siemens Medical Solutions, Erlangen, Germany). Between-group comparison of brain-tissue-corrected levels of these markers (LCModel, SPM12, MATLAB 2023a) and correlation analyses were performed (SPSS v26).

**Results:** Compared with controls, patients showed significantly lower N-acetylaspartate levels (by 14.5%, *p* = *0.03*) but no significant alterations in glutamate-glutamine levels (by -4.0%, *p* = *0.6*). Significant correlations were found between N-acetylaspartate and the glutamate-glutamine complex in patients (*p* = *0.02*), but not in controls (*p* = *0.06*). No significant correlations were found between S1 markers and hand impairment (*p* > *0.05* for all).

**Conclusions:** Our findings demonstrate neuronal mitochondrial dysfunction, which correlates with lower cortical excitability in the ipsilesional S1 in chronic stroke survivors. Further work on the functional relevance of such findings, using larger sample sizes, is warranted.

## Introduction

Recovery of hand motor function after a stroke involves a dynamic process that engages brain areas beyond the motor system^1^. One of these areas is the primary somatosensory cortex (S1), which is well known for its involvement in sensorimotor integration and feedback mechanisms, supporting the planning, execution, and learning of motor skills ^2-6^. Despite this, the last two decades of stroke rehabilitation research have focused on motor remapping or reorganization, particularly of the primary motor cortex (M1, a critical cortical area responsible for hand motor control^7,8^ and recovery^1,9^), leaving S1 remapping understudied (see review^10^). Few studies demonstrated the crucial role of sensorimotor interactions in hand rehabilitation after a unilateral stroke. For example, in the early weeks after stroke, there is a shift in S1 activation (as measured by functional MRI) toward the contralesional, or non-injured, hemisphere^11,12^. This early functional reorganization is followed by a switch back to, or “normalization” of, S1 activation in the ipsilesional, or injured, hemisphere, a process that parallels recovery of skilled hand movements^11^. Non-invasive S1 stimulation also promotes recovery of hand movements in all phases of stroke^13,14^. Briefly, repetitive transcranial magnetic stimulation over the S1, paired with motor practice or sensory stimulation, is likely to enhance learning and recovery of hand movements, thereby improving functional independence. Likewise, sensory stimulation alone appears to increase corticospinal excitability and enlarge hand motor representation, ultimately promoting hand recovery (see review^15^). Despite growing evidence of S1’s role in the rehabilitation of hand movements, no study has investigated whether its metabolic status is altered and how these alterations may contribute to hand impairment or recovery after a stroke. Such understanding may lead to the identification of new targets for restorative therapies focused on tackling a major contributor to disability after stroke, the hand impairment^16^.

Our group was the first to characterize the nature, evolution, and functional relevance of MR Spectroscopy (MRS)-detected metabolic alterations across motor cortices in different phases of stroke^17-23^. Specifically, in the ipsilesional radiologically normal-appearing M1, we found lower levels of the putative marker of neuronal integrity, N-acetylaspartate (NAA), supporting the likelihood of widespread neuronal metabolic downregulation in remote areas connected to the injury site, so-called diaschisis^24-26^. Changes in MRS-detected markers of the balance between excitatory neurotransmission and glial recycling pathways^27^, including glutamate and glutamine measured together as Glx, further support this phenomenon: lower Glx levels in this area suggest a dysregulation of excitatory signaling in M1 controlling the paretic hand. Notably, these alterations are functionally relevant, i) relating to stroke chronicity and severity^17,19^, ii) predicting hand motor outcomes with interventions^21^, and iii) normalizing when hand movements recover ^22^. We also found altered correlational patterns between these two markers compared to those in the healthy population; a higher, positive, and significant correlation was observed between NAA and Glx^19^, indicating a strong relationship between neuronal metabolic status and excitability of that area. This, together with our finding of a significant correlation between NAA levels and the spatial extent of M1 activation during a task performed with the paretic hand^20^, sheds light on the potential neural mechanisms underlying the well-known ipsilesional M1 hypoexcitability after a stroke^28^. Whether similar alterations in the levels and/or correlations of these metabolites occur in ipsilesional spared S1 and how these alterations relate to hand impairment are unknown.

We hypothesized that participants with chronic subcortical stroke would exhibit altered levels of NAA and Glx in the ipsilesional S1, as these alterations have been observed in the ipsilesional M1 in a similar population^17^, and given the extensive anatomical connections between S1 and M1 ^29-31^. We further hypothesized that metabolite correlations would be altered similarly to those observed in M ^17^, with NAA and Glx highly correlated. However, given differences in cellularity and function (see the review^10^), a different magnitude of these alterations could be expected. We also hypothesized that S1 metabolite levels would be functionally relevant, showing a significant relationship to the clinically assessed hand motor impairment.

## Materials and Methods

### Participants

Fourteen chronic stroke survivors and ten age- and sex-matched healthy controls provided informed consent in accordance with the University of Kansas Medical Center Institutional Review Board. Stroke participants were included if they: (i) had experienced a single ischemic stroke at the subcortical level resulting in unilateral hemiparesis at least one month before participation, (ii) had no lesion in S1, thalamus, or brainstem on T2-weighted MRI, (iii) had normal sensation (as detected by Semmes-Weinstein monofilaments^32^), and (iv) sufficient motor function to complete a handgrip task (Fugl-Meyer Upper Extremity Scale^33^, FMUE, greater than 10) since these patients participated to a motor intervention. The data reported here were collected at the baseline (before the intervention). Participants with contraindications to MRI or comorbid neurological, psychiatric, or orthopedic conditions were excluded from the study. Ten healthy controls, free of neurologic, psychiatric, or systemic disorders and demonstrating normal structural MRI findings, were included in the analysis.

### Experimental Protocol

#### Hand impairment evaluation

The FMUE Scale (normal = 66) quantified upper limb motor impairment. Hand strength was assessed bilaterally using a Jamar dynamometer (Asimow Engineering Co., Los Angeles, CA), and the strength of the paretic hand was expressed as a percentage of the non-paretic hand strength (normal > 89%).

#### MRS imaging and analysis

Neuroimaging data were collected using a 3T Siemens MRI scanner. During scanning, the heads of all participants were immobilized with head cushions and instructed not to move. A high-resolution T1-weighted anatomical scan (MPRAGE; TR = 2300 ms; TE = 3 ms; matrix = 256 × 256; voxel size = 1 × 1 × 1 mm^3^) was acquired to guide MRS slab placement and estimate brain tissue composition within the spectroscopic voxels. Axial T2-weighted and proton density images (TR = 4800 ms; TE1/TE2 = 18/106 ms; slice thickness = 5 mm, no gap) were used to verify that the S1 remained radiologically intact and confirm the subcortical stroke location. MRS imaging was performed using a point-resolved spectroscopy (PRESS) sequence (TE = 30 ms; TR = 1500 ms; voxel size = 15 × 15 × 15 mm^3^; spectral width = 1200 Hz). Based on anatomical landmarks, a multivoxel slab was placed across the ipsilesional sensorimotor cortices. Eight 20 mm outer volume suppression bands were used to minimize lipid contamination. Automated, followed by manual, shimming was performed to achieve an optimal full width at half maximum of <20 Hz for the water signal from the entire excitation volume. Motion correction was performed using a rigid-body transformation, estimating six parameters: three translational and three rotational. These parameters were inspected for head movement. None of the participants moved their head more than 2 mm in any direction. The total scan duration was about 45 minutes.

MRS analysis methods have been detailed previously^17^ and consisted of MRS data (LCModel, a linear combination of model spectra using a basis set included in the package, and a radio-frequency coil loading factor^34^) and T1-weighted image analysis (brain segmentation, SPM12, Welcome Department of Cognitive Neurology, London, UK). Three spectroscopic voxels with the following characteristics, gray matter > 75%, a signal-to-noise ratio > 10, and Cramer-Rao lower bounds < 20%, were selected in the hand territory of ipsilesional S1. Using a custom-designed software (Matlab 2023a), each voxel’s metabolite values were corrected for partial volume effects ^35^ using the formula: c = cLCModel × [1/Fbrain], where Fbrain = % grey matter + % white matter within the spectroscopic voxel, derived from brain tissue segmentation in SPM12 (Wellcome Department of Cognitive Neurology, London, UK). The levels of each metabolite in the selected voxels were averaged and expressed in millimoles per kilogram of wet weight brain tissue. Since more than half of the patients had left hemisphere injury (57%; **Table 1**), for those with the stroke located in the right hemisphere (n = 6), we right-to-left flipped the data so that the “left” hemisphere was the ipsilesional one. This flip was performed based on the lack of significant differences between hemispheres in MRS-detected metabolites in the healthy population^17,18^. Thus, the ipsilesional S1 metabolites were compared with those in the left S1 in controls.

**Table 1.** Demographic and clinical data for stroke participants (n = 14), including age, sex (M, male; F, female), time since stroke (months, mo), affected hemisphere (L, left; R, right), stroke location, and Fugl-Meyer Upper Extremity (FMUE, normal = 66) scores and handgrip strength (expressed as a percentage of the non-paretic hand, normal > 89%) at recruitment.

| Age/Sex | Time post-onset<br>(mo) | Site of stroke | FMUE | Hand<br>strength (%) |
| --- | --- | --- | --- | --- |
| 57/M | 6 | L / basal ganglia, corona radiata | 66 | 91.5 |
| 48/F | 1 | R / corona radiata | 66 | 91.2 |
| 71/F | 5 | R / corona radiata | 66 | 89.3 |
| 68/M | 2 | L / posterior insula | 66 | 91.6 |
| 75/M | 1 | L / internal capsule | 62 | 98.3 |
| 65/M | 3 | R / basal ganglia | 62 | 90.3 |
| 48/F | 11 | L / internal capsule, basal ganglia | 61 | 93.6 |
| 45/F | 27 | R / basal ganglia, corona radiata | 36 | 41.9 |
| 61/M | 15 | L/Basal ganglia | 30 | 30.8 |
| 64/M | 30 | R / basal ganglia, internal capsule,<br>corona radiata | 27 | 3.7 |
| 66/F | 2 | L / internal capsule | 26 | 8.8 |
| 59/M | 144 | R / internal capsule, basal ganglia | 25 | 10.3 |
| 58/M | 27 | L / internal capsule, basal ganglia | 24 | 8.6 |
| 61/M | 24 | L / basal ganglia, internal capsule,<br>corona radiata | 10 | 7.2 |

#### Statistical Analysis

Statistical analyses were performed using a combination of parametric and non-parametric tests, depending on data distribution (as determined by the Kolmogorov-Smirnov test). If data were non-normally distributed, a paired-sample Wilcoxon signed-rank test and Spearman correlation test were used. If normally distributed, a paired t-test and Pearson correlation test were used. Group comparisons and correlation analyses were conducted between stroke participants and age, sex, and school-year-matched healthy controls. All analyses used SPSS (v26.0), and results were two-tailed with α = 0.05. For visualization, we used the % change in the averaged metabolite levels between the stroke and control groups (% change NAA = [(NAA stroke - NAA control) / NAA control] * 100).

## Results

### Participants’ characteristics

Stroke and control participants did not significantly differ in age (mean ± SD = 60.4 ± 8.8 vs. 54.0 ± 10.8 years, *p = 0*.*1*), sex distribution (35.7% vs. 40.0% women, *p = 0*.*4*), or years of education (14.3 ± 2.0 vs. 14.9 ± 2.5; *p = 0*.*6*). All patients had sustained a single subcortical or brainstem infarction (in one patient) for at least one month before testing (**Table 1**) and had radiologically intact S1 on T2-weighted imaging. Patients demonstrated a wide range of arm/hand impairment (FMUE = 44.8 ± 20.9, from 10 to 66) and hand strength (54.1% ± 40.9% of the non-paretic side, from 3.7% to 98.3%, **Table 1**).

### Spectra quality was high in both groups

Signal-to-noise ratios for all spectra in each group were within recommended tolerances (11.7 ± 1.4 in stroke participants, from LCModel vs. 12.7 ± 1.6 in controls, *p = 0*.*1*; **Fig. 1A**). The full width at half maximum of metabolites included in analysis was about 12 Hz in all participants, reflecting high spectral resolution [13].

**Fig. 1.**
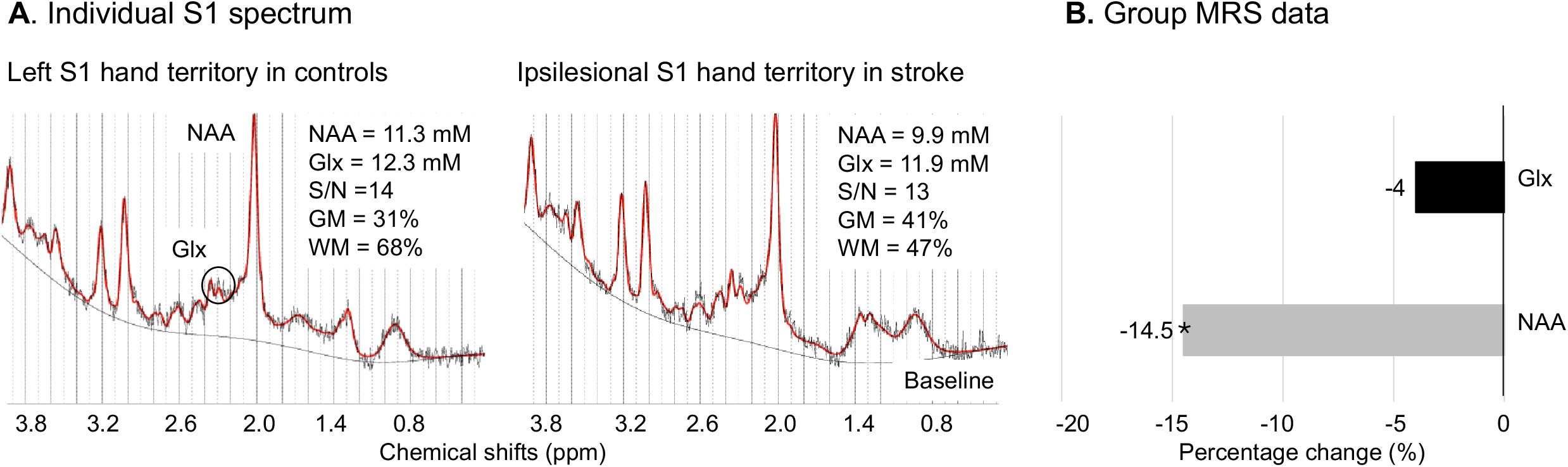
**(A)** LCModel output from spectroscopic voxels in the left (in a control participant) and ipsilesional (in a patient) S1 hand territories. The output consisted of the acquired MRS spectrum (black line), estimated spectrum (red line), and underlying baseline. Spectrum quality was high, with a signal-to-noise (S/N) of 14 for the control and 13 for the patient. Metabolite concentrations were calculated (LCModel) and corrected for the brain tissue composition within the voxel (grey matter, GM + white matter, WM). **(B)** Percentage changes in NAA (grey column) and Glx (black) in stroke participants versus controls showed significantly lower NAA levels (*, p<0.05).

### The brain tissue composition in the spectroscopic voxels was similar between the groups

Similar S1 brain tissue (grey matter plus white matter) compositions were found between groups (84.0 ± 10.5% vs. 85.1 ± 7.3%, *p* = *0.8*).

### Stroke participants exhibit lower NAA levels but unaltered Glx levels

As predicted, patients showed significantly lower NAA levels than controls (mean = 9.6 ± 1.0 vs. 11.2 ± 2.1 mM; z = - 2.2, *p* = *0.03*; change = -14.5%). This result was specific to NAA and was not found for Glx (10.4 ± 2.9 vs. 10.9 ± 1.8 mM; z = 0.5, *p = 0*.*6*; change = -4.0%, **Fig. 1B**). We also examined the effect of time after stroke onset, sex, and age at diagnosis on metabolite levels; we found no significant effects, with *p-values* ranging from *0*.*09* to *0*.*7*.

### Stroke participants exhibited significant metabolite correlations

We found significant correlations between NAA and Glx in stroke participants (r = 0.60, *p = 0*.*03*), but not in controls (r = 0.61, *p = 0*.*06*).

### S1 metabolites did not correlate with hand motor impairment

Contrary to our prediction, we failed to detect statistically significant correlations between NAA or Glx levels and FMUE or hand strength (*p-values* ranged from *0*.*1* to *0*.*9*).

## Discussion

To our knowledge, this is the first study to provide evidence of alterations in neuronal integrity in the radiologically normal-appearing S1 in patients with chronic stroke. As predicted, compared to controls, stroke participants exhibited lower levels of NAA, a putative biomarker of neuronal integrity, in the somatosensory hand territory located contralateral to the paretic hand. NAA levels were further positively associated with Glx levels, suggesting that metabolically depressed neurons are likely to exhibit lower excitability. Contrary to our predictions, we failed to find significant between-group differences in Glx levels nor significant correlations between NAA or Glx and hand impairment. Below, we discuss these findings in detail, along with their implications and the limitations of our study.

### Lower levels of NAA are found in the S1 hand territory contralateral to the paretic hand

Consistent with our predictions, we found lower NAA levels in the S1 hand territory compared to the analogous territory in age- and sex-matched healthy controls (see **Fig. 1B**). The magnitude of NAA decrease (by 14.5%) is similar to those reported in the hand territories of the ipsilesional M1 (lower by 14.2%^17^) or supplementary motor cortex (13.9%^18^) in a similar population. Consequently, we consider the differences in NAA levels reported here, relative to those in controls, to be reliable and likely to reflect a similar phenomenon observed in the motor cortices. Lower levels of NAA are thought to indicate neuronal metabolic downregulation and/or neuronal death (see review^36^). Because the MRS voxels were selected in radiologically intact S1 distant to injury, and no damage to the thalamus or brainstem was detected, the observed NAA alterations are unlikely to be indicative of neuronal death resulting from overt structural damage or neurodegeneration. Moreover, given that similar brain tissue composition exists within the selected spectroscopic voxels, neuronal death also appears unlikely. Alternatively, lower NAA could be due to decreased cerebral blood flow, likely caused by carotid stenosis, which may be present in these patients (and was not evaluated in this study). Such blood flow alteration would be expected to have a more global effect on NAA levels. Since we didn’t find altered NAA levels in the ipsilesional dorsal premotor cortex in a very similar population^18^, we speculate that altered blood flow is unlikely to explain our NAA findings. Therefore, lower NAA may instead reflect neuronal metabolic downregulation, likely due to diaschisis^24-26^. Briefly, diaschisis refers to the suppression of neuronal activity, metabolism, or perfusion in spared brain regions remote from, but connected to, the injury site, resulting from disruption of afferent or efferent pathways. Yet, diaschisis may also contribute to a range of long-term cellular-level metabolic alterations, such as abnormalities in mitochondrial function or decreased metabolic activity; lower NAA supports these changes. This phenomenon could also lead to long-term morphological changes, i.e., shrinkage in body cell size, reduction in the number of synaptic buttons, incomplete endings, and loss of synapses; we did not find significant differences in grey/white matter volume between patients and controls. However, we cannot exclude microscopic neuronal changes. Finally, diaschisis could lead to changes in the excitability-inhibition balance; we found non-significant changes in Glx levels in S1 (see detailed discussion below). Still, the positive and significant correlation between NAA and Glx levels indicates a direct relationship between neuronal metabolic status and excitability level. A similar enhanced NAA-Glx coupling was observed in the ipsilesional M1^17^, further supporting our interpretation.

### Glx levels in S1 hand territory contralateral to the paretic hand were not significantly altered

It is important to note that the differences observed between stroke participants and controls were specific to NAA and were not evident for Glx (**Fig. 1B**). The lack of significant Glx alterations could be partially explained by differences in cellularity between S1 and M1. Indeed, approximately half of the synapses in M1 are exclusively excitatory, whereas in S1, there are more synapses formed with both excitatory and inhibitory neurons^37^. Anatomically, S1 is heavily connected with M1, with excitatory projections to M1^29,30^ and mostly inhibitory projections from M1^31^, enabling complex information integration during sensorimotor control and learning. This S1-M1 infrastructure may be shaped during stroke rehabilitation, and such reshaping may contribute to our findings. Yet, how this infrastructure is impacted by stroke rehabilitation is unknown.

### S1 NAA and Glx levels did not correlate with clinically detected hand impairment

Contrary to our prediction, we did not find significant correlations between NAA or Glx levels and clinical impairment. The relatively wide time range post-stroke (from one month to 12 years) may mask the role of S1 neuronal and excitability changes in recovery from the impairment, since initially suppressed ipsilesional S1 excitability increased over time^11,12^. However, the nature of these relationships is unclear: Do these neuronal-level changes somehow contribute to the hand impairment, or do they have a reduced sensitivity to motor performance, or are they responses to chronic altered afferent information to S1 or altered M1-S1 connectivity? Further research is necessary to fully understand the nature of these relationships.

### The current study has several limitations

The first limitation is the sample size; although comparable to prior MRS studies in this population, it limits statistical power and may have prevented the detection of subtle effects. Second, the variance of time since stroke is relatively large. Because we examined patients from one month to more than a decade after stroke, we have no hint from the present data of the possible time course of S1 NAA or Glx changes.

## Conclusion

In summary, we found altered neuronal integrity in the spared S1 hand territory years after a unilateral subcortical stroke. This finding, along with the robust coupling between neuronal integrity and cortical excitability, provides a means to assess the fine-grained underpinnings of possible diaschisis in S1. Understanding how stroke impacts sensory information processing and/or flow will help design effective therapeutics for hand motor impairment.

